# Montane rain forest dynamics under changes in climate and human impact during the past millennia in northern Madagascar

**DOI:** 10.1101/2023.06.07.544115

**Authors:** Vincent Montade, Laurent Bremond, Helena Teixeira, Thomas Kasper, Gerhard Daut, Elysée Rasoamanana, Perle Ramavovolona, Charly Favier, Fabien Arnaud, Ute Radespiel, Hermann Behling

## Abstract

Madagascar comprises one of the Earth’s biologically richest, but also one of most endangered, terrestrial ecoregions. Although it is obvious that humans substantially altered its natural ecosystems during the past decades, the timing of arrival of humans on Madagascar as well as their environmental impact is not well resolved. In this context, this research aims to study and compare the influence of early human impact and climate change on rain forests and wildlife in northern Madagascar during the past Millennia. By using palaeoenvironmental reconstructions from lake sediment cores in a montane environment (Montagne d’Ambre), results indicate a major drought, starting approximately 1,100 years ago. This drought caused significant changes in lake levels and vegetation dynamics. Human impact, evidenced by fires, started a few decades later. Anthropogenic burning, limited to the low-altitude areas, was therefore not the driving force behind these early changes observed in the lake catchment areas. Although this does not dismiss the strong impacts humans had subsequently on these ecosystems, this work demonstrates that the late Holocene natural drought that intensified regionally about one thousand years ago, significantly impacted the ecosystems independently and prior to anthropogenic activities. At a regional scale, a review of demographic studies revealed a substantial number of inferred population bottlenecks in various wildlife species during the last millennia, likely resulting from this combination of both human-related impact and natural environmental changes (i.e., precipitation decline). This research highlights that the current state of ecosystems in northern Madagascar results from both human impact and natural climate changes. It also points to the importance of a multi-site and multi-proxy comparison for deciphering the nature and succession of past environmental changes.

## 1. Introduction

Tropical rain forests are essential to humankind, for goods, services, carbon sequestration, and therefore, for the balance of global biogeochemical cycles [1]. However, a large part of rain forests is facing a rapid increase of anthropogenic pressures mainly through deforestation [2]. Deforestation is often linked to the expansion of human populations and is frequently the result of the development of agricultural practices carried out at the expense of forests.

Madagascar represents a typical example; fast-human population growth coincided with accelerated forest loss and fragmentation over the past decades [3]. Most of the remaining rain forests are scattered across eastern Madagascar and in mountainous areas containing small ‘pockets’ of rain forests. Since Madagascar is a major hotspot of biodiversity [4], these ’pockets’ are critical for protecting the unique biodiversity and in particular forest-dependent taxa, but also for securing the water resources of the island. At higher elevations, air masses generate frequent fogs and high orographic precipitation [5]. This moisture is recycled through evapotranspiration processes of rain forests which also prevent soils from runoff and erosion and thus contribute greatly to total water discharges for the surrounding lowlands. Some of the steeper mountain areas in Madagascar still remain relatively unaffected by human impacts, while on their foothills (at lower altitudes) human impacts are generally stronger. This discrepancy may be explained partly by the official protection of certain mountain areas but also by some climatic constraints (i.e., high rainfall and cool temperatures) and steep slopes slowing down the settling of human populations. For assessing the resilience of such forests, it is critical to study their responses to environmental changes with present-day datasets and over long timescales that go beyond historical records, and to analyse a broad range of ecological trajectories. However, obtaining high resolution palaeoecological records in such environments encompassing several millennia is difficult, as it requires sedimentary records from perennial humid areas (e.g., lake, peat bog, marsh) which are not frequent in the tropics.

In Madagascar, a first study has been realised in such an isolated mountain rain forest in the northern part of the island in the National Park (NP) of Montagne d’Ambre (Figure 1 and [6]). This NP represents a highly suitable area for such an approach, as several crater lakes occur at different elevations within the same mountainous area (Figure 1). The first sediment archive, taken from a lake located at 1,250 m above sea level (asl) near the mountaintop, has enabled the reconstruction of the main vegetation changes during the past 25,000 years before present (BP). Vegetation records have revealed a strong influence of the African Humid Period (AHP) in northern Madagascar, which shaped this study site with a replacement of montane forest by evergreen humid forest. The AHP ended at around 5,500 yr BP, marked by a precipitation decline, as evidenced in East Africa [7]. Over the last millennium, a peat bog developed at this study site which currently covers a large part of the lake. Simultaneously, fires are starting to be recorded more frequently and seem to reflect the beginning of human impacts surrounding Mt. d’Ambre NP. This result is consistent with archaeological data showing an increase of human activities recorded in different regions mainly around the last millennium [8]. On the other hand, increase of anthropogenic activities hardly explains the development of the palustrine and peat bog vegetation, which should rather be the result of the filling up of the sedimentary basin and/or the result of an hydrological change. A gold standard in palaeoecology for disentangling drivers of past environmental change on vegetation dynamics is to provide multi-site reconstructions.

**Figure 1.**
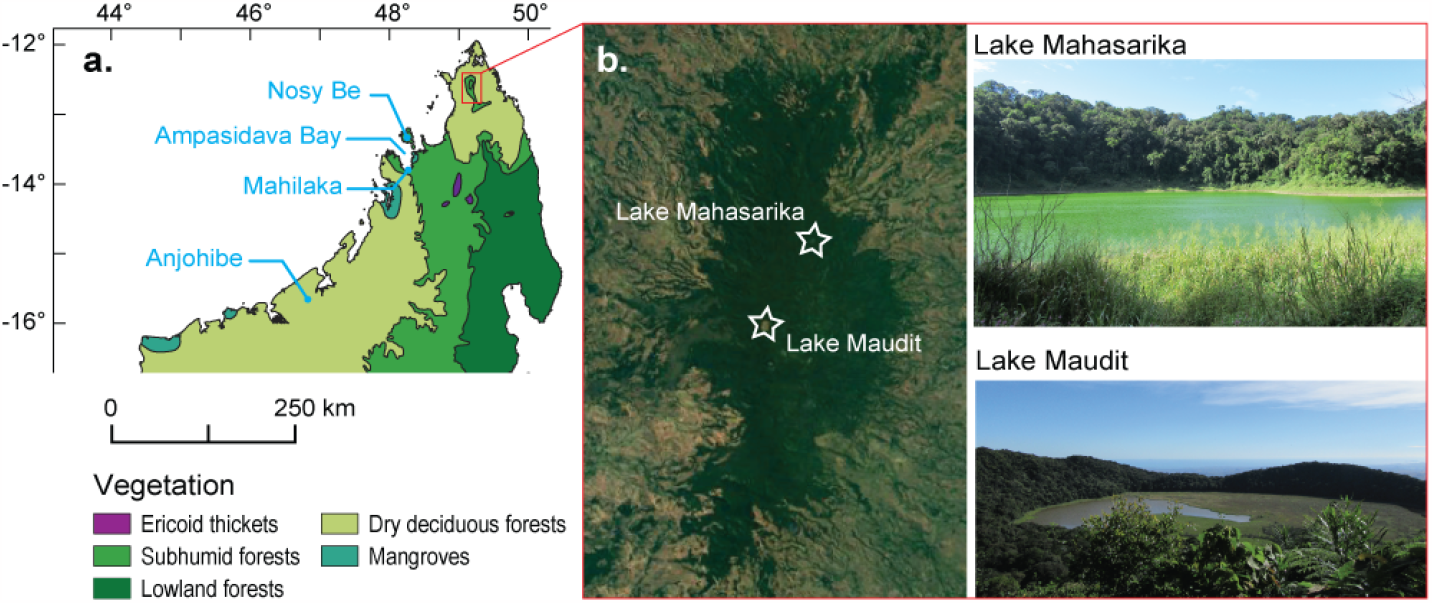
a Distribution map of natural biomes in Madagascar [9]. b Satellite image of Montagne d’Ambre (source: Google Earth) indicating location of Lake Mahasarika and Lake Maudit.

In this context, our study of a new lake site at a lower elevation in Mt. d’Ambre (Lake Mahasarika,1,073 m asl) aims to better understand and infer the regional responses, and not purely site-related responses, of such a ’pocket’ of rain forest to past environmental changes. More specifically, based on multi-site comparison, our detailed analysis of late Holocene (the past 4,000 yr BP) changes aims to disentangle potential different drivers of environmental and forest dynamics. Additionally, the palaeoecological results will be compared with our present knowledge on corresponding demographic dynamics of wildlife from northern Madagascar.

These new data from Mt. d’Ambre and comparisons with available data will contribute to answer the following questions: When did humans start to impact ecosystems in northern Madagascar? To what extent have rain forests and wildlife been impacted locally and regionally? Is there evidence for natural climate changes impacting vegetation changes over the last millennia?

## 2. Material and methods

### (a) Palaeoenvironmental study site

Montagne d’Ambre NP is a volcanic massif reaching up 1,475 m asl, that spreads 35 km north-south and 15 km east-west located in northern Madagascar at about 10 km south of Antsiranana. In this region, the climate is characterised by heavy rainfalls in austral summer, which are related to the intertropical convergence zone in northern Madagascar when it moves to its southern boundary [10]. The study site, Lake Mahasarika, is one of the six crater lakes distributed at different elevations on Mt. d’Ambre. Surrounded by a dense humid montane rain forest, Lake Mahasarika is located on the north-eastern flank of the massif at an elevation of 1,073 m asl. Due to its higher elevation, climate conditions in Mt. d’Ambre contrast with surrounding lowlands. Air cooling with increasing altitude generates frequent fogs and important orographic rainfalls (annual precipitation at “La Station Forestière de la Roussette” reaches above 3,000 mm.yr^-1^ [11]) allowing the development of a dense rain forest which contrasts to the dry forest, savanna and pastures situated in the lowland areas surrounding the mountain. This small isolated patch of forest on the mountain is essential for the local human population in terms of water resources (maintaining soil, etc.) and can be expected to be very sensitive to past precipitation changes. Lake Mahasarika has been selected for its shallow depth (< 10 m) allowing the use of light sediment coring materials on a lake, for its good accessibility, and for its proximity to the ecotone between humid forest and sub-humid/dry forest at about 800 m asl [12]. Although the sub-humid/dry forest has almost entirely disappeared near to the park mainly due to human impact over the past decades [3], the vegetation dynamics surrounding the lake may at least partially capture the dynamic ecotone changes, and certainly better so than forest located at higher elevations.

### (b) Sediment coring and age-depth modelling

Fieldwork was carried out in August 2018. Based on bathymetry, the sediment core LMAHA-18 was taken from the central and deepest part of the lake (7 m water depth; 12.53509S, 49.17701E). Core LMAHA-18 was sampled with a Livingstone piston sediment corer operated from a small platform settled on two rubber boats. Using sampling tubes of ca. 5 cm diameter and 120 cm length, a total of 6 sections of about 100 cm sediment length were sampled with overlaps of 10-50 cm length. Preserved in aluminium tubes for transport to the laboratory, sediment was extruded from each aluminium tube and split in two sister cores, LMAHA-18a and LMAHA-18b. Each sister core was stored under cool (4°C) and dark conditions at the laboratory facilities (LMAHA-18a at the Friedrich-Schiller-University Jena and LMAHA-18b at the University of Goettingen, Germany). Cores were photographed and magnetic susceptibility was scanned in 3 mm steps with three replicate measurements using a MS2E surface scanning sensor (Bartington Instruments). Using lithological description, patterns of magnetic susceptibility, and specific marker layers (electronic supplementary material, Figure S1), the different core sections were aligned to a composite master sequence LMAHA-18. The age-depth model is based on 16 AMS radiocarbon dates of 15 bulk sediment samples and 1 plant macro-remain (Table S1), performed at either the Poznań radiocarbon laboratory (Poland) or the LMC14 (Gif-sur-Yvette, France) laboratory. The age of the sediment surface is considered as modern, and was thus set to the year of coring (AD 2018 = - 68 calibrated yr BP). The age model was performed as a function of the composite depth with the RStudio software using the R-package “Bacon” (V. 4.0.5) [13] and using SHCal20 calibration curve and bomb ^14^C curve [14,15].

### (c) Palaeoecological analyses

For pollen extraction, 28 subsamples of 0.5 cm^3^ from LMAHA-18 were processed following standard chemical techniques with chloridric acid, potassium hydroxide, fluorhydric acid and acetylosis [16]. One tablet of exotic *Lycopodium* spores (20848 +/- 1546, batch number 1031) was added to each sample to estimate pollen concentration. A minimum sum of 300 terrestrial pollen grains was counted for each subsample using a light microscope at 400x magnification, and pollen and fern-spore percentages were calculated on the terrestrial pollen sum. Pollen and spore identification were based on several atlases [17–20], the online African Pollen Database (https://africanpollendatabase.ipsl.fr/#/home) and the reference collections from University of Goettingen (http://www.gdvh.uni-goettingen.de/) and the University of Montpellier (https://data.oreme.org/observation/pollen). To investigate compositional changes in the tree community, a Principal Component Analysis (PCA) was performed on square-root transformed relative abundances of arboreal pollen taxa, defined as the percent count of each taxon relative to the total number of arboreal pollen grains in a sample (electronic supplementary material, Figure S3). For extraction of charcoal particles, 1 cm^3^ of sediment was taken every cm along the LMAHA-18 core. Each sample was soaked in a 3% NaP_2_O_4_ solution plus bleach for several hours to deflocculate the sediment and oxidise organic matter. The samples were sieved through a 160 μm mesh and the carbon particles were counted using a x40 magnification stereomicroscope. Bulk organic δ^13^C was measured against certified standards (L-Prolin, EDTA and USG65) and reported in standard δ notation (‰) against Vienna Pee Dee Belemnite (VPDB) and Air, respectively. Relative errors based on triplicate measurements are 0.05 ‰ for δ^13^C.

### (d) Demographic dynamics of wildlife in northern Madagascar

A comprehensive review was conducted on published molecular population demographic studies conducted on Malagasy taxa from northern Madagascar. Studies were collected from the literature using a combination of the following keywords: population genetics, population dynamics, phylogeography, demography, demographic modelling, coalescence, bottleneck, expansion, Quaternary climate change, vegetational shifts and Madagascar. All studies were carefully screened to confirm that the study species were in fact distributed and sampled in northern Madagascar, roughly defined as the area ranging from Nosy Be to the Loky–Manambato region, to achieve the best possible congruence with the palaeoenvironmental records. Studies on species that only occurred outside northern Madagascar were excluded from our review. We reviewed the evidence from studies using different molecular markers (i.e., mtDNA, microsatellite loci and genome-wide SNPs) and various demographic approaches. Studies that fulfilled our criteria were evaluated regarding taxonomic representation (i.e., mammal, bird, amphibian, non-avian reptile, plants) and habitat type (i.e., adapted to dry forest, humid forest or both habitats. Finally, the direction of the detected population demographic change (i.e., constant population size, population bottleneck, expansion or size recovery) was assigned to the palaeoecological periods of interest whenever possible. When multiple studies were available for a single taxon, they were all included in the review for comparative reasons.

## Results

Combining pollen, charcoal, sedimentological changes and isotopic data, three distinct periods were distinguished within the Late Holocene which were each characterised by different plant assemblages.

### (a) Environmental conditions from 4,000 years to 1,100 yr BP (Period 1)

During this period the sediment was dark to light brown, i.e., homogeneous and organic rich. Except for some peaks of magnetic susceptibility, interpreted as event related deposits (electronic supplementary material, Figure S1), the sediment accumulation rate (SAR) was relatively low (0.1 cm/yr) and remained constant showing low erosion.

With a relatively low proportion of Poaceae (around 10%, Figure 2b), the rain forest was well-established around the study site, as it is also indicated by the dominance of arboreal pollen (*Mallotus, Noronhia, Celtis* and Moraceae-Urticaceae; electronic supplementary material, Figure S2 and Table S1). However, the slow decline in PCA Axis-1 values (Figure 2a) reflects that changes in tree composition have occurred during this period. Such changes are marked by, e.g., *Noronhia* peaks (24%) at ca. 2,550 yr BP. *Mallotus* increased progressively and reached a maximum (27%) at ca. 1,150 yr BP (electronic supplementary material, Figure S2). *Mallotus* is a potential indicator of early successional pioneer trees and its increase may provide evidence for increased disturbances. Low values of δ^13^C generally below -24 ‰ also confirm the dominance of C3 forest vegetation in the catchment of the lake (Figure 2d). Furthermore, few charcoal particles are recorded only occasionally, indicating a very low level of fire activity (Figure 2e and 2f).

**Figure 2.**
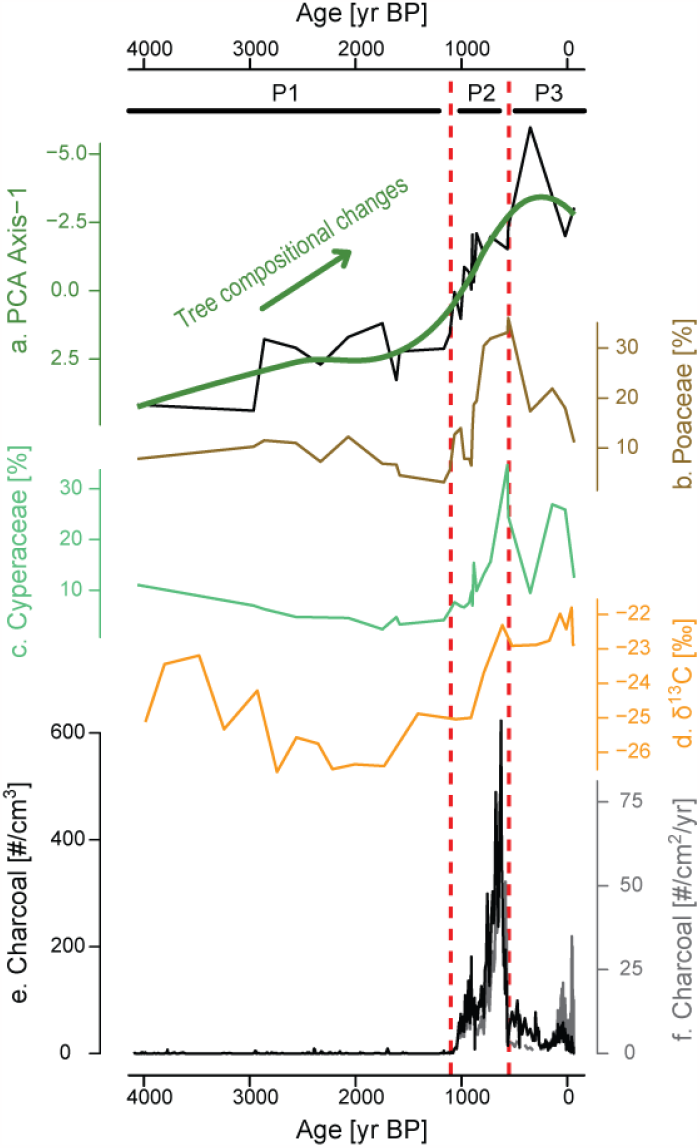
Synthetic palaeoenvironmental reconstructions obtained from the sediment core recovered in Lake Mahasarika (MAHA-18). a Axis-1 of the Principal Component Analysis performed on arboreal pollen taxa reflecting tree compositional changes. b Poaceae pollen grains. c Cyperaceae pollen grains. d δ^13^C measured on bulk organic fraction. e Concentration (black) and f influx (grey) of charcoal particles (>160 μm). The vertical lines demarcate the succession of the three main periods (P) of different environmental conditions.

### (b) Ecological changes and fire increase between 1,100 and 550 yr BP (Period 2)

During this period, the sediment had the same texture as in the previous period and SAR remained broadly in the same range with values slightly higher between 0.1 and 0.2 cm/yr (electronic supplementary material, Figure S1). On the other hand, a major vegetation change was recorded and characterised by a two-step increase of Poaceae and Cyperaceae pollen from 1,100 and 900 yr BP up to 30% each (Figure 2b and 2c). The abrupt decrease in the values of axis-1 of the PCA indicates a major change in the tree composition from 1,100 yr BP onwards (Figure 2a). This shift was slightly preceded by a change in forest dynamics (electronic supplementary material, Figure S2), with a synchronous decline of *Mallotus* (<10%) and increase of *Trema* (>10%). *Trema* is generally associated with light demanding trees living at forest edges [12]. It may have outcompeted *Mallotus* with increased disturbances right before the increase of herbs. From 900 yr BP, the changes of the tree composition continued (Figure 2a). Arboreal pollen taxa related to low disturbances and humid rain forest habitat decreased (e.g. *Celtis, Podocarpus, Pandanus*) at the expanse of new tree taxa (*Garcinia, Ziziphus, Ilex*, Araliaceae) showing that a tipping point was reached and the rain forest composition changed substantially (electronic supplementary material, Figure S2 and Table S2). The presence of *Impatiens* (electronic supplementary material, Figure S2), typical of small shrubs growing in disturbed rain forest and in the rain forest margins, also provides evidence for an increase of forest disturbance. As pollination of *Impatiens* is entomophilous (producing less pollen than anemophilous plants), pollen grains of *Impatiens* have a low dispersal probability. These pollen grains were more likely deposited near the parent plants, indicating that disturbances occurred locally at the study site. The δ^13^C values around -22‰ from 900 yr BP (Figure 2d) indicate an increased proportion of C4 vegetation within the catchment of the lake. This confirms that vegetation changes (increase of Poaceae and Cyperaceae) recorded by pollen data occurred locally at the study site. Charcoal particles also showed a two-step increase in values (Figure 2e and 2f). A first increase occurred from ca. 1,070 yr BP onwards and reached rapidly values above 70 particles/cm^3^ and 10 particles/cm^2^/yr with a peak around 900 yr BP. The second one occurred from 800 yr BP onwards and reached values above 200 particles/cm^3^ and 30 particles/cm^2^/yr with a maximum at around 600 particles/cm^3^ and 100 particles/cm^2^/yr at ca. 630 yr BP. The increase in fires might be responsible for increased forest disturbances and development of herbs. If fires would have occurred in the catchment area of the lake, forest fires would have resulted in strong sediment discharges into the lake, since it is characterised by a small catchment area (ca. 2.3 km^2^) and steep slopes. However, the increase of fires was not combined with synchronous increase of magnetic susceptibility showing no major increase of erosion from 1,100 yr BP onward (Figure 4). Consequently, fires certainly occurred in the region of Mt. d’Ambre but not locally at the lake catchment and do not well explain local vegetation dynamics evidenced by pollen and δ^13^C data (Figure 2d). Another potential explanation of local increase of forest disturbances could be a decrease of the lake level related to a decrease in precipitation. Indeed, an important lake level decrease may have allowed a development of C4-grass vegetation (Poaceae) at the shore of the lake and Cyperaceae in shallow lake margins as it is observed today. This lake level decrease, resulting from reduced precipitation, would also be a possible explanation of tree compositional changes.

### (c) Vegetation and fire dynamics since the last 550 yr BP (Period 3)

After 550 yr BP, sediments turned to light brown and SAR started to increase rapidly from 250 yr BP, reaching values above 0.2 cm/yr at ca. 100 yr BP, and then values above 1 cm/yr during the past century (electronic supplementary material, Figure S1). Near the core top, the SAR is expected to increase as the sediment compaction is reduced. Among the main vegetation changes, values of Poaceae pollen decreased progressively from 30 to 10% and Cyperaceae fluctuated between 25 and 10% (Figure 2b and c). Except in one sample, values of PCA axis-1 remain in the same range (Figure 2a), showing that the new tree composition was maintained with the dominance of *Mallotus, Elaeocarpus, Norhonia, Trema* and Araliaceae, between 4 and 10%, respectively (electronic supplementary material, Figure S1 and Table S2). The taxa occurring in the previous period (*Ziziphus, Gacinia, Ilex*), were still represented with the same range of values between 2 and 4%. The δ^13^C values remained around -22‰, showing a mixture of C3 and C4 plants within the catchment of the lake (Figure 2d). This confirms the relatively high proportion of herbaceous vegetation after 550 yr BP. Although charcoal particles were found in every cm of the core (Figure 2e and 2f), the concentration and influx showed a general decrease with values generally below 50 particles/cm^3^ and 10 particles/cm^2^/yr. During the last century, charcoal influx increased due to SAR increase with values above 10 particles/cm^2^/yr and with maximum values around 30 particles/cm^2^/yr while charcoal concentration remained in the same range of values.

### (d) Demographic dynamics of wildlife in northern Madagascar

A total of 17 demographic studies covering 19 studied species distributed across northern Madagascar fulfilled our criteria and were considered here (electronic supplementary material, Table S3). The studies were biased towards mammals (47.4%) and non-avian reptiles (36.8%), with the remaining taxa (amphibian, birds and plants) being represented by a single species each (5.3%). For non-avian reptile species, the timing of the demographic event is either unknown (*Phelsuma dorsivittata*) or dated back to the late Pleistocene (>12,000 yr BP), precluding conclusions about recent demographic dynamics. Studies on mammals, however, provide important insights about recent demographic dynamics in Malagasy wildlife. Except for three species for which demographic events were not dated (*Microcebus tavaratra* [21], *Eliurus tanala* [22] and *Microgale brevicaudata* [23]), all other studied terrestrial mammals (i.e., lemurs and rodents) suffered a population decline predating or during the late Holocene (i.e., before or during Period 3; Figure 3 and electronic supplementary material, Table S3). A reduction in population size was also reported for *Chaerephon leucogaster* [24], a bat species, during the Late Holocene (Period 3 ;but see [25]). Notably, demographic modelling with the composite-likelihood method implemented in fastsimcoal2 for *Microcebus arnholdi* [6] revealed two consecutive population bottlenecks, one predating Period 3 (∼5,000 yr BP) and one during Period 2 (∼1,000 yr BP). Demographic modelling with the Approximate Bayesian Computation (ABC) for two additional lemur species (i.e., *Propithecus perrieri* and *P. tattersalli* [26]) also detected a population decline within the last millennium (Period 2, Figure 3).

**Figure 3.**
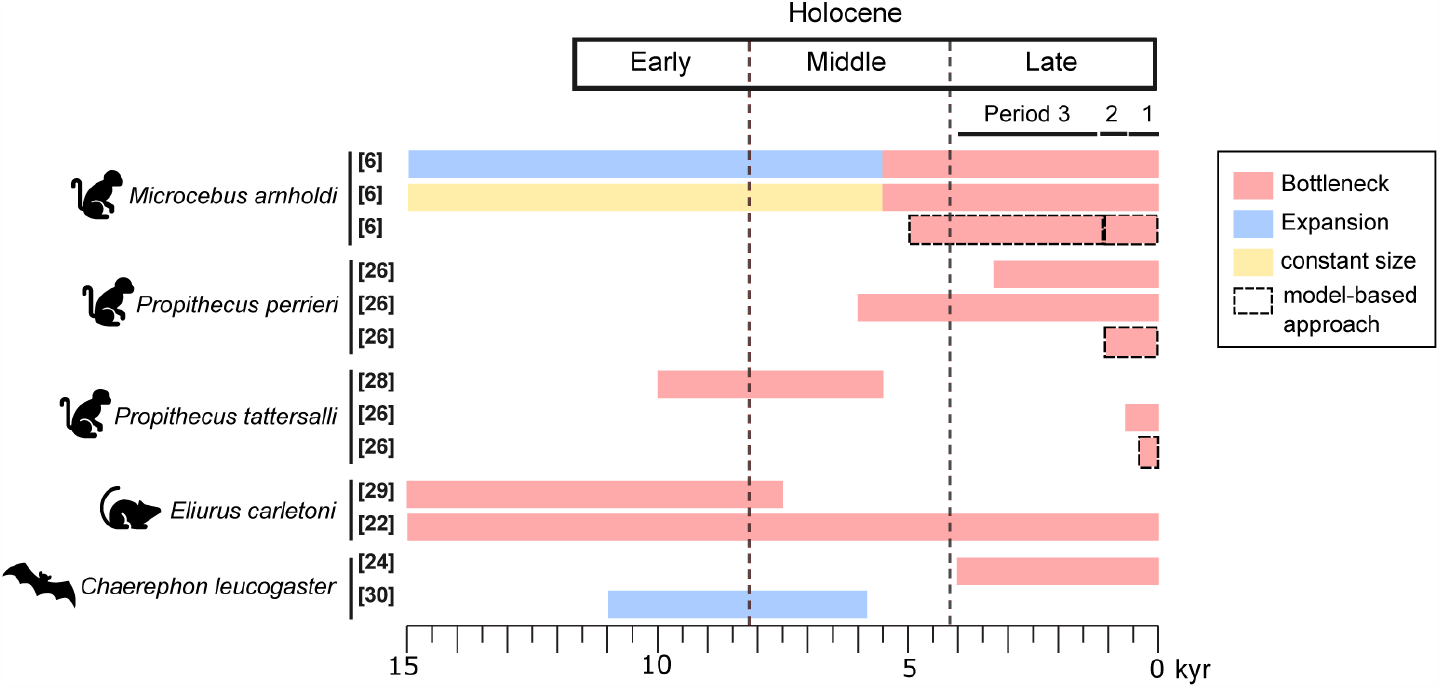
Review results on population demographic dynamics during the past 15,000 years for Malagasy wildlife occurring in northern Madagascar based on molecular datasets. Only studies with dated demographic events are included in the figure (see electronic supplementary material Table S2 for a review of all the studies currently available). The red, blue and yellow bars represent distinct demographic dynamics (population bottleneck, expansion and constant size, respectively). Dashed bars highlight methods that use model-constrained approaches (i.e., *fastsimcoal2* and *ABC*). The remaining studies rely on model-free methods. The late Holocene was divided in the three periods highlighted by the palaeoenvironmental reconstructions of Lake Mahasarika and defined in the text [6,22,24,26,28–30].

**Figure 4.**
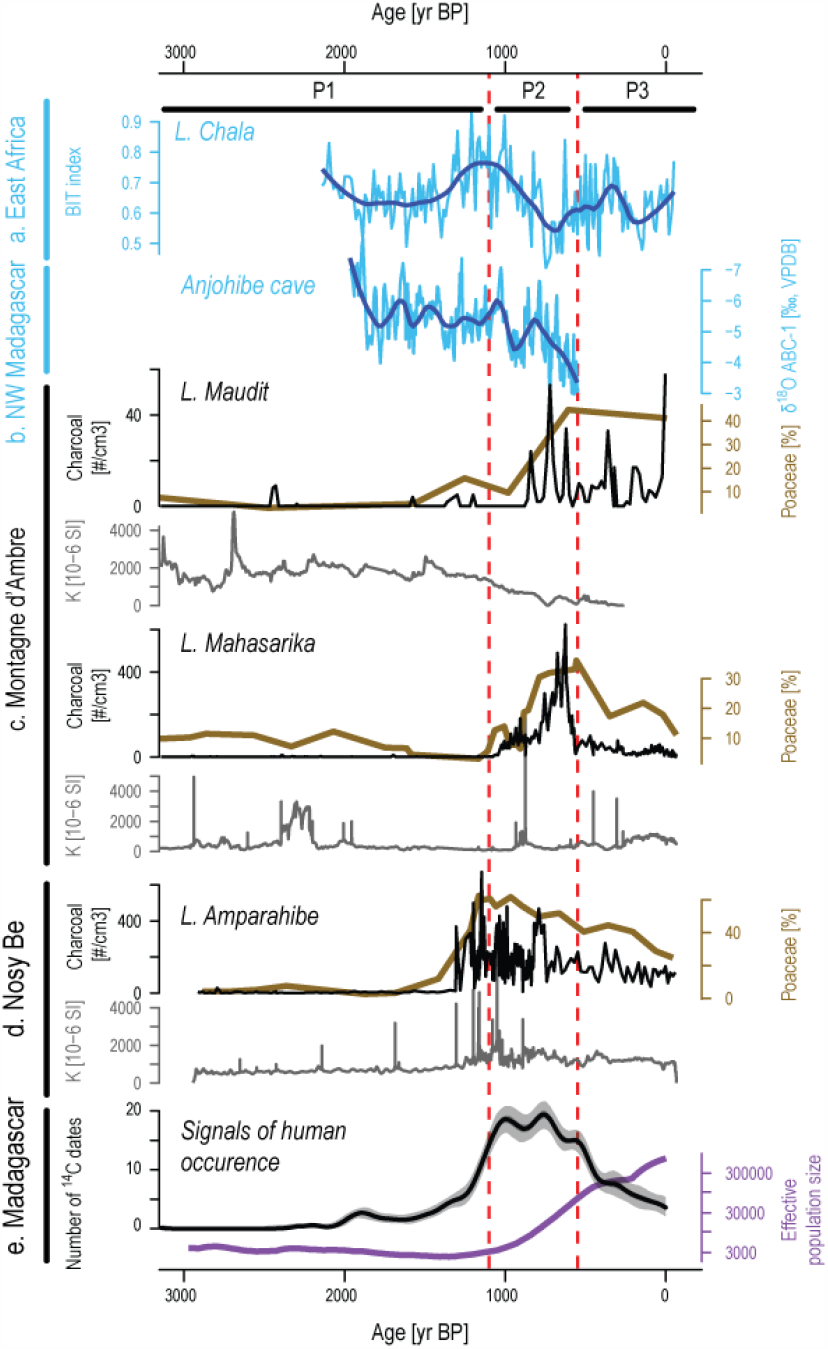
Synthesis of environmental changes during the past 3,000 yr BP. a and b Decadal-resolution time series of BIT index variability from Lake Challa [39] and δ^18^O speleothem profile from Anjohibe Cave [10] showing precipitation changes in East Africa and Northwest Madagascar (the dark blue line represents a smooth spline). c and d Comparison of Poaceae percentages, charcoal concentrations and magnetic susceptibility (K) between Lake Maudit [6], L. Maharika (this study) from Northern Madagascar and L. Amparahibe [37] from Northwest Madagascar. e Black curve represents the number of radiocarbon dates performed on archaeological samples obtained from an archaeological data synthesis [8]. Purple curve represents the estimated changes in the effective human population size [40].

Species demographic response to past environmental vs. human pressures may differ among ecological specialist vs. generalist taxa. The reviewed studies revealed that all mammals adapted to dry habitats exhibited signatures of a population decline (*Propithecus perrieri, Eliurus carletoni, Chaerephon leucogaster*, Table S3). Except for *Furcifer pardalis* (that underwent a population expansion in the late Quaternary; [27]) all taxa adapted to humid habitats also underwent a population bottleneck or kept a constant population size (*Microcebus arnholdi, Eliurus tanala, Mantella crocea, Phelsuma dorsivittata*). In contrast, no clear demographic trend was evident in species adapted to both dry and humid habitats (*Microcebus tavaratra, Propithecus tattersalli, Microgale brevicaudata, Myotis goudoti, Dicrurus forficatus, Laliostoma labrosum, Calumma boettgeri, Geckolepis maculata, Mimophis occultus, Zonosaurus madagascariensis*), suggesting that ecological specialised taxa are more prone to environmental changes than generalist taxa.

Except for [26] and [6], all studies assumed that population structure was negligible, and applied either a single demographic approach (statistical tests or a model-free approach) or a Bayesian Skyline Plot (BSP) coupled with mismatch distribution/ neutrality tests. For *Microcebus tavaratr*a [21] and *Eliurus tanala* [22], the neutrality tests detected a population expansion, while BSP suggested a constant population size. The neutrality tests were also inconclusive for *Eliurus carletoni* [22]. In contrast, the demographic methods (i.e., model-free and model-constrained approaches) implemented by [26] and [6] revealed a synchronous population bottleneck for *Microcebus arnholdi, Propithecus perrieri* and *P. tattersalli* during the late Holocene (period 2 and 1).

## Discussion

### (a) The period prior to permanent human settlement in northern Madagascar

The exact schedule for the settlement of humans in Madagascar is still partly debated [e.g., 8,31,32]. Recent evidence for early to mid-Holocene (>5000 yr BP) arrival of humans on the island is solely based on indirect indicators (e.g., cut-marks performed on hippopotamus and extinct elephant birds bones [31]), and no archaeological artefacts have so far been found associated with these findings. This suggests that even if humans were already inhabiting Madagascar during these early times, they seem to not have impacted the ecosystems sufficiently enough to leave traces in the existing palaeoecological records. The clear first signals of permanent human settlements on Madagascar are generally available for the time period between 2,000 and 1,000 yr BP [e.g., 8,33,34]. However, the earliest proof of human civilization with town establishment in Madagascar has been dated at 1,000 yr BP and it stems from the harbour of Mahilaka in the Northwest. Located at the end of a bay, Mahilaka was inhabited by muslim population and was connected to the Comoro Islands and probably east Africa by maritime trade routes [35,36]. This population seems to be related to the westward Austronesian expansion that had developed maritime trade and agricultural practices based on rice cultivation in the region from 1,250 yr BP onwards [36]. Correspondingly, human impact with an increase of anthropogenic burning has been recently suggested on the near-by Malagasy Nosy Be Island from 1,300 yr BP onwards [37]. Our new data from Lake Mahasarika combined with previously published data from Lake Maudit show very low occurrences of fires prior to 1,100 yr BP (Figure 4). This suggests that permanent human settlements did not exist in the areas surrounding the Mt. d’Ambre before this time and that the observed vegetation dynamics would thus have been caused by natural factors such as climate changes.

Palaeoclimate records from Mid-to Late Holocene from northwestern Madagascar have revealed a multi-millennial drying trend in this region combined with frequent occurrences of megadroughts [10,38]. Sedimentological changes at higher altitude of Lake Maudit (1,250 m asl) in Mt. d’Ambre in Northern Madagascar are consistent with an overall drying trend that intensified at the end of the AHP from 5,500 yr BP onwards [6]. Starting from 4,000 yr BP, Mahasarika pollen assemblages reveal that dominant tree taxa were continuously changing (Figure 2) and *Mallotus* progressively increased until 1,100 yr BP. As a typical taxon of early successional trees, it might indicate that rain forest, adapted to more drought-related disturbances or mesic conditions, may have developed due to the effect of reduced rainfall and occurrences of megadroughts. Forest taxa from Lake Maudit did not show such changes, suggesting that forest was not affected by climate change at higher altitudes [6]. As humidity increases with elevation in Mt. d’Ambre due to decreasing temperatures, a regional decline in precipitation should have affected the vegetation at lower altitudes, such as at Lake Mahasarika (1,073 m asl), more than rain forests near the mountain top, such as at Lake Maudit (1,250 m asl), where orographic precipitation and cloud cover is maximum. This altitude-dependent effect seems to be confirmed by our study which suggests that ecosystems from lower elevations may be more affected by regional precipitation decline than mountain environments which are under the influence of orographic rainfall that still maintains rain forest. A palaeoecological record from an even lower elevation would allow us to test this hypothesis for this region. The available demographic studies for Northern Madagascar showed that all mammals, birds, amphibians, and non-avian reptiles ranging from Nosy Be to the Loky–Manambato region exhibited a signature of a population size change (expansion or decline) pre-dating the Late Holocene, supporting major environmental changes in the region long before humans impacted these ecosystems.

### (b) Human impact and environmental changes in Northern Madagascar

Since 1,100 yr BP, the permanent occurrence of charcoal particles with increase of influx combined with the development of grasses in the Mahasarika record suggests a fire activity increase related to an increase of anthropogenic burning in the surrounding lowland region. On Nosy Be Island, charcoal particles increased strongly together with grasses and were correlated with peaks of erosion, supporting local disturbances in the catchment from anthropogenic burning and this about 200 years earlier than in Mt. d’Ambre (Figure 4). However, the pattern of changes evidenced from the Mahasarika record differ in more than just dating. First, the tree composition changes that suggest increased disturbances with a maximum of *Mallotus* followed by an increase of *Trema*, a tree typically growing in disturbed forests and at forest edges, slightly preceded the fire increase. This shows that the fire increase detected by the Mahasarika charcoal record was preceded by increased disturbances and not vice versa. Second, the increase of charcoal particles was not associated with a marked peak of erosion which excludes local fires at the lake catchment. Consequently, the local development of grasses and sedges evidenced by pollen assemblages and δ^13^C results with the establishment of new forest composition cannot be explained by the increase of fire frequency within the lake catchment. By contrast, a lake level decrease, allowing the development of herbs and sedges (reaching maxima between 900 and 550 yr BP) at the lake shore, as it can be seen today, would explain better the observed results from Lake Mahasarika. The complementary results from Lake Maudit suggest the same overall process; the charcoal increase was also not associated with any increase of local erosion during the last millennium (Figure 4). Erosion signal in the central part of the lake (coring site) was even reduced by the development of a peat bog from ca. 1,000 yr BP onwards, also suggesting a lake level decrease at that time combined with the local development of grasses and sedges. Altogether, local changes evidenced in Mt. d’Ambre are initially (ca. 1,100 - 1,000 yr BP) better explained by natural precipitation decline and lake level decline, and then by the increase of anthropogenic burning near Mt. d’Ambre. Major precipitation decreases or a megadrought, starting around 1,000 yr BP and culminating at ca. 900 yr BP, is also supported by palaeoclimatic proxies from other sites in Madagascar [10,38,41,42], the Island of Rodrigues [10] and East Africa [39]. The Mt. d’Ambre palaeoecological records reveal that the increase of anthropogenic burning and human expansion occurred during the subsequent period of reduced humidity, contemporaneous of the Medieval Climate Anomaly in East Africa and in the Northern Atlantic region (ca. 1,000-700 yr BP [43,44]). A combination of natural and anthropogenic factors may subsequently have led to the dramatic ecological changes observed in Northern Madagascar, such as additional animal population declines evidenced by demographic modelling for *Microcebus arnholdi* (Mt. d’Ambre NP), *Propithecus perrieri* (Analamerana – Andrafiamena) and *P. tattersalli* (Loky – Manambato; reviewed here) or megafauna extirpation, as revealed by archaeological data [45,46]. Although a wide range of approaches are currently available and were used to reconstruct species demographic history from molecular data (e.g., neutrality tests, model-free and model-based approaches), the inference of recent population bottlenecks during the last millennia was only successful with model-based approaches, such as fastsimcoal2 and ABC. Moreover, while some studies, implementing mismatch distribution/neutrality tests and Bayesian Skyline Plots, detected different demographic trends, the studies implementing model-free and model-based approaches [6,26] revealed congruent dynamics. Altogether, our demographic review confirms that model-based approaches are a very valuable tool to detect recent demographic changes that are not detected by other demographic approaches, and highlights the importance of using more than one method for demographic inferences.

Change in subsistence strategies with populations shifting from hunter-gatherer to herding-farming have also been proposed as a potential explanation of the major ecological changes around 1,000 yr BP [47–50]. However, considering the strong hydroclimatic change during this period, and without additional palaeorecords combined with archaeological data to obtain a robust regional synthesis, the causal links between the observed changes, in particular during the critical period of transition, cannot be fully clarified at the island scale. In particular it remains open, if anthropogenic burning increased because precipitation decreased which may also have led human population to use and develop new substance strategies and/or colonise new areas. It should also be considered whether the precipitation decrease may have promoted anthropogenic fire expansion and the occurrence of disasters such as megafires.

### (c) Period of the last 550 yr BP

Over the past few centuries, data from Lake Maudit have not shown particular changes and vegetation dynamics but instead mainly reflected the development of a large local peat bog [6]. In contrast, from 550 yr BP onwards at Lake Mahasarika and, starting from ca. 200 years earlier at Nosy Be, the proportion of grasses decreased in congruence with a decrease of fires. This pattern, particularly the decrease of fires, might be explained in several ways: First, starting approx. 600 yr BP, archaeological data suggested a human population decline in the Northwest at Mahilaka and the surrounding villages of Ampasindava Bay [51,52]. Although this remains to be confirmed at the local and regional scale around Mt. d’Ambre with archaeological data, a decline in human occupation in Northern Madagascar could have resulted in a reduction of anthropogenic burning. Second, several precipitation records have revealed an increase in humidity from about 500 yr BP onwards [41]. This may have reduced the risk of catastrophic fire events, such as megafires, and may also explain a general reduction in fires. Third, in case Northern Madagascar was entirely forested, the abrupt increase of fires at a regional scale at the beginning of the last millennium might have produced large amounts of charcoal particles. Subsequently, and as a consequence of more frequent fires, fuel availability for large fires should have decreased, which could also explain the observed decrease in carbon particles emitted. These different hypotheses, or a combination of several of them, seem likely but require additional archaeological and palaeoecological data to be tested. Finally, during the last century, a renewed increase of fires is recorded in the Mahasarika sediment core (Figure 2e and f). Additional study sites are necessary to confirm this trend which is evidenced by charcoal influx. A fire increase during the last century might reflect the effect of the European colonisation in the Mt. d’Ambre region, where Jofreville has been established with development of agriculture from the 1900s [53].

## 5. Summary

This multi-proxy and multi-site comparison of lacustrine cores, located in the North of Madagascar in the mountain range of the Mt. d’Ambre NP, revealed a major drought, starting approximately 1,100 years ago, causing a significant drop in the water level at the two lacustrine study sites (Lakes Mahasarika and Maudit). Under low lake levels, palustrine and peat bog vegetation started to develop on the shore of the two lakes. In addition, the rain forest closer to the edge of the montane rain forest zone has been affected by precipitation decrease, as indicated by an increase of pioneer trees. Recorded a few decades later, anthropogenic burning spread throughout the study region, but was limited to the low-altitude areas surrounding the Mt. d’Ambre. Very likely fires were therefore not the driving force behind these early changes observed in the lake catchment areas, as it has also been observed for other study sites, such as on the island of Nosy Be [37]. Although this does not dismiss the importance humans had subsequently in terms of impact on ecosystems, this work demonstrates that the natural drought that intensified regionally about one thousand years ago, significantly impacted the ecosystems independently of anthropogenic activities. The increasing number of inferred population bottlenecks of wildlife at a regional scale during the last millennium, as evidenced by the review of demographic studies, likely resulted from the combination of both human-related impact and environmental changes (i.e., precipitation decline). Remaining important questions, therefore, are not just how precisely humans have influenced their environments, but also whether the natural precipitation decrease caused humans to change their substance strategies and/or to move to new areas. Although the current increase in anthropogenic activities is eroding the resilience of forests, these montane rain forests have shown substantial resilience to climate change during the past millennia. Continuing to protect such areas therefore seems very relevant and essential in the context of global warming. Finally, this work shows that multi-proxy approaches and multi-site comparisons provide essential evidence for distinguishing the different factors driving environmental changes in order to better understand ecosystem functioning over the long term.

## Supporting information

Electronic supplementary material

## Acknowledgements

We thank Madagascar National Parks, the director of Mt. d’Ambre National Park, the Direction du Système des Aires Protégées, the Direction Générale du Ministère de l’Environnement et des Forêts de Madagascar, Madagascar’s for their permission to conduct the field work in Mt. d’Ambre National Park (N°120/18/MEEF/SG/DGF/DSAP/SCB.Re) and for their support. We thank Romule Rakotondravony for his support during the palaeoecological fieldwork. We thank the guides of Mt. d’Ambre National Park and the local communities for their indispensable help in the field. Five ^14^C dates were acquired with the help of Tomasz Goslar (Poznan Radiocarbon Laboratory). Ten ^14^C analyses were acquired thanks to the CNRS-INSU ARTEMIS national radiocarbon AMS measurement programme at Laboratoire de Mesure ^14^C (LMC14) in the CEA Institute at Saclay (French Atomic Energy Commission). This study was funded by the Deutsche Forschungsgemeinschaft (RA 502/20-1, RA 502/20-3, and BE 2161/30-1) and PALEOMAD project (INSU-LEFE/CNRS).

