## Supplementary material for "Montane rain forest dynamics under changes in climate and human impact during the past millennia in northern Madagascar": Electronic supplementary material

#### Sediment core description and chronology from Lake Mahasarika

Core LMAHA18 is characterised by a brown to dark brown organic-rich clay sediment with various greyish to reddish intercalated distinct layers from 1.5 to 6 cm thick. Magnetic susceptibility fluctuates with values around 300 $10^{-6}$ SI and shows various peaks with values above 2000 $10^{-6}$ SI generally matching well with the greyish to reddish distinct layers. Changes of magnetic susceptibility, colours and occurrence of distinct layers allowed to identify the different overlaps between the six sediment core sections. All overlaps have been confirmed except one between section 3 and 4. Between these two sections, ca. 10 cm of sediment is missing (Figure S1). Following this core parallelization, the composite master core LMAHA18 has a total length of 475 cm and can be divided into five main stratigraphic units.

From depth 475 to 375 cm (unit 1), the sediments are laminated and contain five distinct layers. Thickness of lamination varies between 0.3 and 3 cm and colours alternate between brown and dark brown. Laminations are no longer visible from the second unit (unit 2) between 375 and 335 cm depth. In this unit sediment colour remains dark brown and only one distinct layer is recorded. From the third unit (unit 3), the sediment lightens slightly and turns brownish at 335 cm until 300 cm depth. The magnetic susceptibility values are higher than in other units (between 1000 and 2000 $10^{-6}$ SI) and three distinct layers are visible. In the fourth unit (unit 4), between 300 and 115 cm depth, the colour turns back to dark brown and eleven distinct layers are identified. Of the eleven distinct layers, one records the maximum peak of magnetic susceptibility with values reaching 9000 $10^{-6}$ SI. In the last unit (unit 5), from 115 cm to the top core, the sediment colour is homogeneous and changes to light brown. No distinct layers are intercalated in this unit.

The distinct layers, corresponding to greyish to reddish intercalated layers generally associated with a peak of magnetic susceptibility, were considered as event-related deposits. These event-related deposits are assumed to be quick, within only hours or maybe days. They were thus omitted from the age-depth modelling which is based on 16 Accelerator Mass Spectrometry radiocarbon dates (Table S1). All the radiocarbon ages are in stratigraphic order and none are reversed. The model has a basal age of 4,100 calibrated years Before Present (yr BP) and the sediment accumulation rate (SAR) varies between 0.04 and 1.6 cm $yr^{-1}$. SAR values fluctuate generally between 0.05 and 0.2 cm $yr^{-1}$ in the first four sedimentological units. Major increases of SAR are recorded in the last unit (unit 5) from 90 cm with values above 0.2 cm $yr^{-1}$ and from 40 cm with values above 1.5  cm $yr^{-1}$.

**Table S1** Accelerator Mass Spectrometry (AMS) radiocarbon ages of composite sediment core LAMAHA18.

| Sample code | Core section | Composite depth [cm] | Material | 14C Age [yr BP] | Error [+/-] | pMC [%] | Error [+/-] | 2σ calibrated age [yr BP] |
|---|---|---|---|---|---|---|---|---|
| SacA62281 | LMAHAa-1 | 43.5 | Bulk | Modern | | 115.43545 | 0.26743 | -41.5 - -40.5 |
| SacA-62282 | LMAHAa-2 | 88.083 | Bulk | 95 | 30 | | | 5 - 251 |
| Sac-A62283 | LMAHAa-2 | 121.083 | Bulk | 255 | 30 | | | 145 - 313 |
| Poz-111278 | LMAHAa-2 | 146.083 | Bulk | 625 | 30 | | | 530 - 644 |
| Sac-A62284 | LMAHAa-3 | 157.789 | Bulk | 685 | 30 | | | 557 - 659 |
| Sac-A62285 | LMAHAa-3 | 181.789 | Bulk | 965 | 30 | | | 766 - 920 |
| Poz-111280 | LMAHAa-3 | 196.789 | Bulk | 1050 | 30 | | | 803 - 958 |

| | | | | | | | |
|---|---|---|---|---|---|---|---|
| Sac-A62286 | LMAHAa-3 | 212.789 | Bulk | 1195 | 30 | | 961 - 1175 |
| Sac-A62287 | LMAHAa-3 | 225.789 | Bulk | 1250 | 30 | | 999 - 1259 |
| Poz-115913 | LMAHAa-4 | 258.5 | Leaf | 1745 | 30 | | 1538 - 1699 |
| Sac-A62288 | LMAHAa-4 | 294.5 | Bulk | 2190 | 30 | | 2016 - 2304 |
| Sac-A62289 | LMAHAa-5 | 352.905 | Bulk | 2485 | 30 | | 2358 - 2704 |
| Poz-110924 | LMAHAa-5 | 402.905 | Wood | 2875 | 30 | | 2853 - 3068 |
| Sac-A62290 | LMAHAa-6 | 442.103 | Bulk | 3565 | 30 | | 3694 - 3955 |
| Poz-107795 | LMAHAa-6 | 472.603 | Bulk | 3755 | 35 | | 3929 - 4224 |

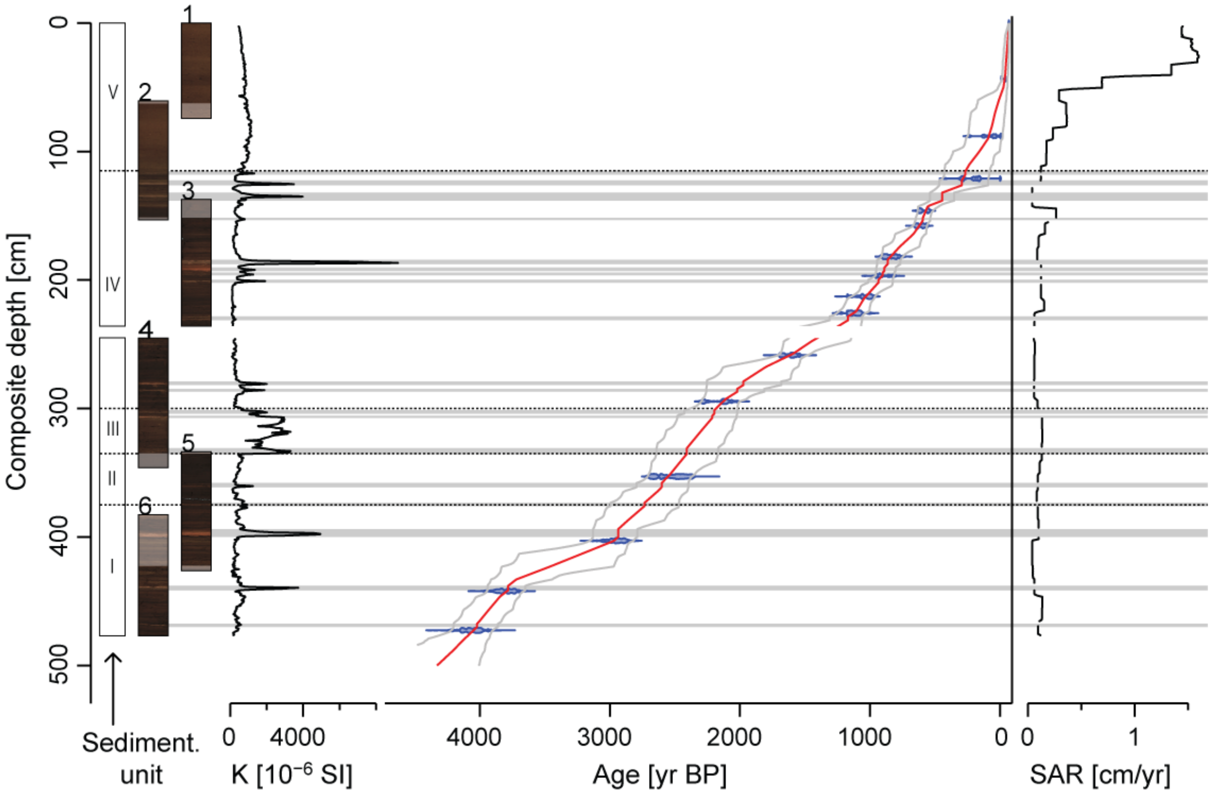


**Figure S1** Sedimentological units, digital images of each sediment core section, magnetic susceptibility (K), age-depth model and sediment accumulation rate (SAR) for the composite sediment core LMAHA18. The composite sediment core was established based on the parallelization of visual marker layers as well as the magnetic susceptibility of the six core sections. Age-depth model is based on a Bayesian model [1] with medians and the 2σ error ranges of the 15 Accelerator Mass Spectrometry (AMS) radiocarbon ages. The AMS radiocarbon ages were calibrated using the SHCal20 dataset [2] and bomb $^{14}C$ curve (zone 1–2) [3].

### Palynological data from Lake Mahasarika

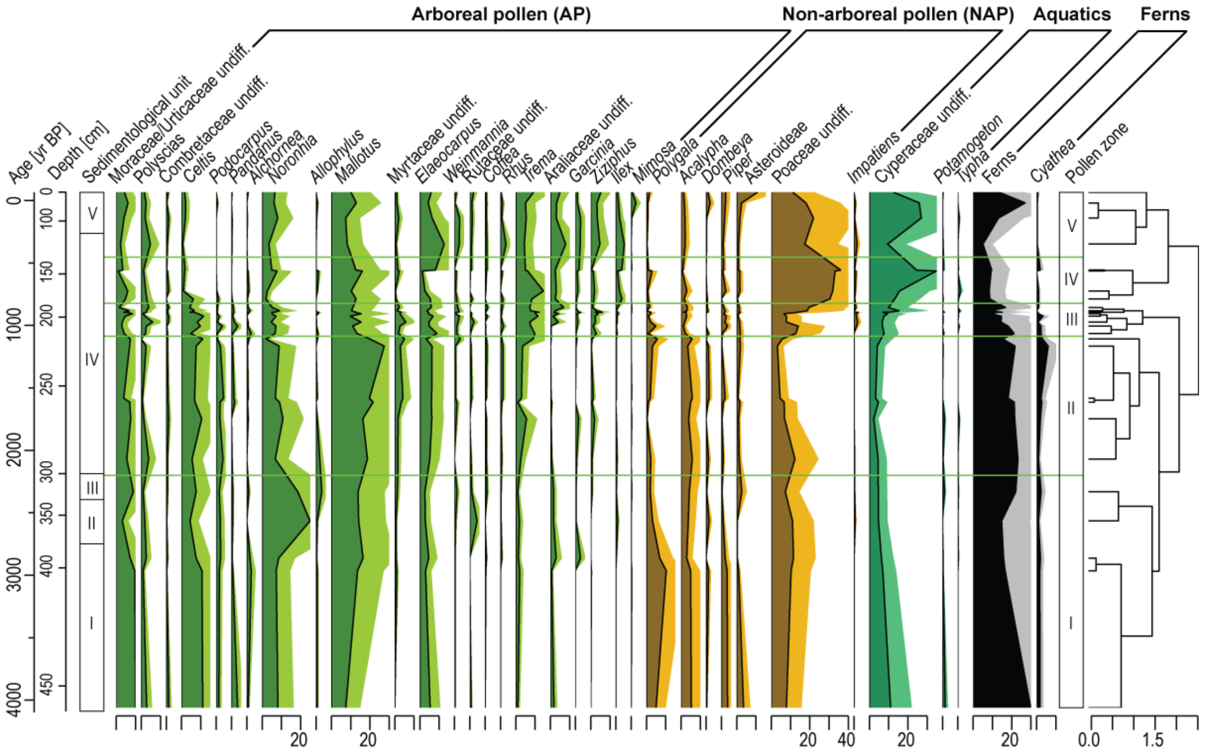


**Figure S2** Pollen and fern spores records of composite sediment core LMAHA18 plotted on age scale (indicated in calendar years before present) and on depth. Percentages of pollen and fern spores were calculated based on the terrestrial pollen sum. Pollen zones indicated by green lines were identified by a cluster analysis (CONISS, [4]). The light coloured curves correspond to exaggerated curves by a factor 5.

**Table S2** Summary of the pollen zones from Lake Mahasarika, indicating their depth and age range and their dominant terrestrial pollen taxa.

| Pollen Zone | Depth range [cm] | Age range [kyr BP] | Dominant terrestrial pollen taxa |
|---|---|---|---|
| PZ5 | 135-0 | 0.5-0 | Poaceae, *Mallotus*, *Elaeocarpus*, *Noronhia*, *Trema*, Araliaceae |
| PZ4 | 180-135 | 0.8-0.5 | Poaceae, *Mallotus*, *Trema*, *Elaeocarpus*, *Noronhia*, Moraceae/Urticaceae |
| PZ3 | 222-180 | 1.1-0.8 | Poaceae, *Mallotus*, *Trema*, *Celtis*, *Noronhia*, *Elaeocarpus* |
| PZ2 | 306-222 | 2.2-1.1 | *Mallotus*, *Celtis*, *Noronhia*, Poaceae, Moraceae/Urticaceae, *Elaeocarpus* |
| PZ1 | 475-306 | 4.1-2.2 | *Noronhia*, *Mallotus*, Poaceae, *Celtis*, Moraceae/Urticaceae, *Polygala* |

**Principal component analysis with arboreal pollen taxa**

In order to condense the information of the arboreal community changes through time, a Principal Component Analysis (PCA) was carried out, using the tree taxa plotted in the synthetic pollen taxa (see Figure S2) as input variables (Figure S3).

The PCA results in one main axis, the axis-1 representing 26% of the total variance. Axis-1 shows high positive values for *Celtis*, *Podocarpus* and Moraceae/Urticaceae and high values loadings for *Ilex, Ziziphus, Garcinia*, Araliaceae and *Trema*. This axis resumes well the main arboreal taxa changes during the past, showing a continuous tree compositional changes over the past 4,000 yr BP and showing a marked increase of tree compositional changes from 1,100 yr BP. The next three axes explain lower values of the total variance, around 10% for each of them. As highlighted for Axis-2 in Figure S3 (not shown for Axis-3 and -4), these axes do not represent any significant pattern reflecting consistent tree community compositional changes. These axes are therefore not considered for the interpretation of tree community changes through time.

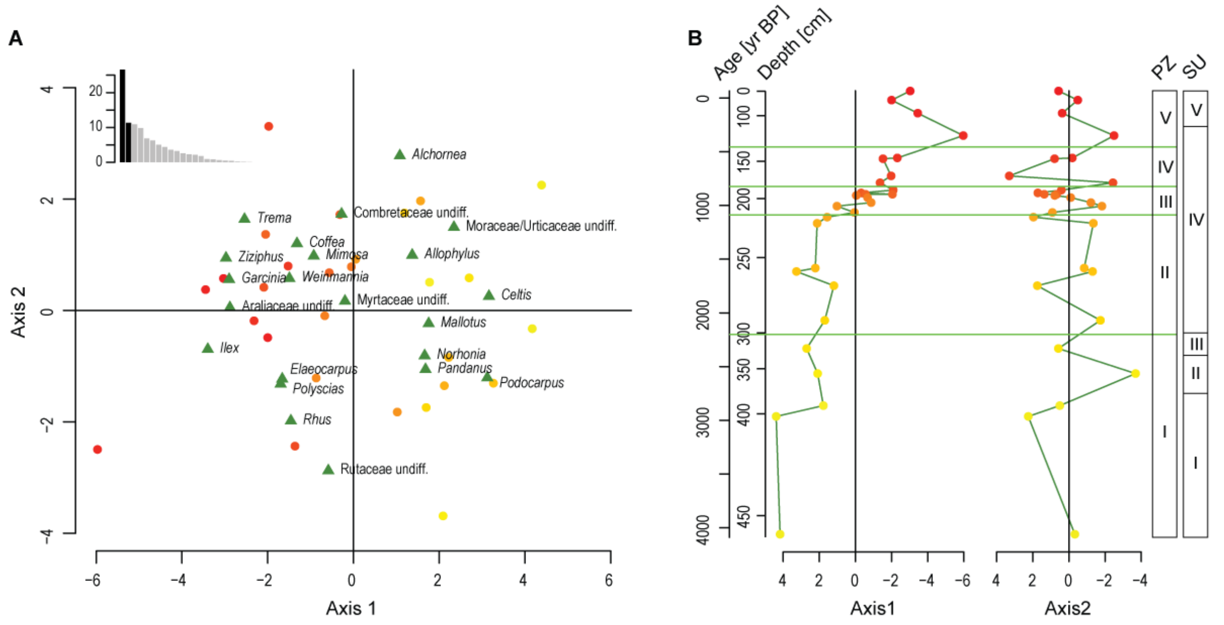


**Figure S3** Principal component analysis (PCA) of Arboreal pollen (AP) taxa of the composite sediment core LMAHA18. A, PCA bi-plot for axis-1 and -2 and histogram of cumulative variances of the PCA axes. Dots indicate the distribution of samples from yellow to red according to their respective ages. Green triangles indicate the distribution of the AP taxa. B, Values of samples for PCA axis-1 and -2 plotted on age scale (indicated in calendar years before present) and on depth. The dots show the same coloured gradient according to respectives ages of samples. The bars on the right side represent the main sedimentological units (SU) and the pollen zones (PZ) also evidenced by the green lines.

**Sedimentary charcoal particles from Lake Mahasarika**

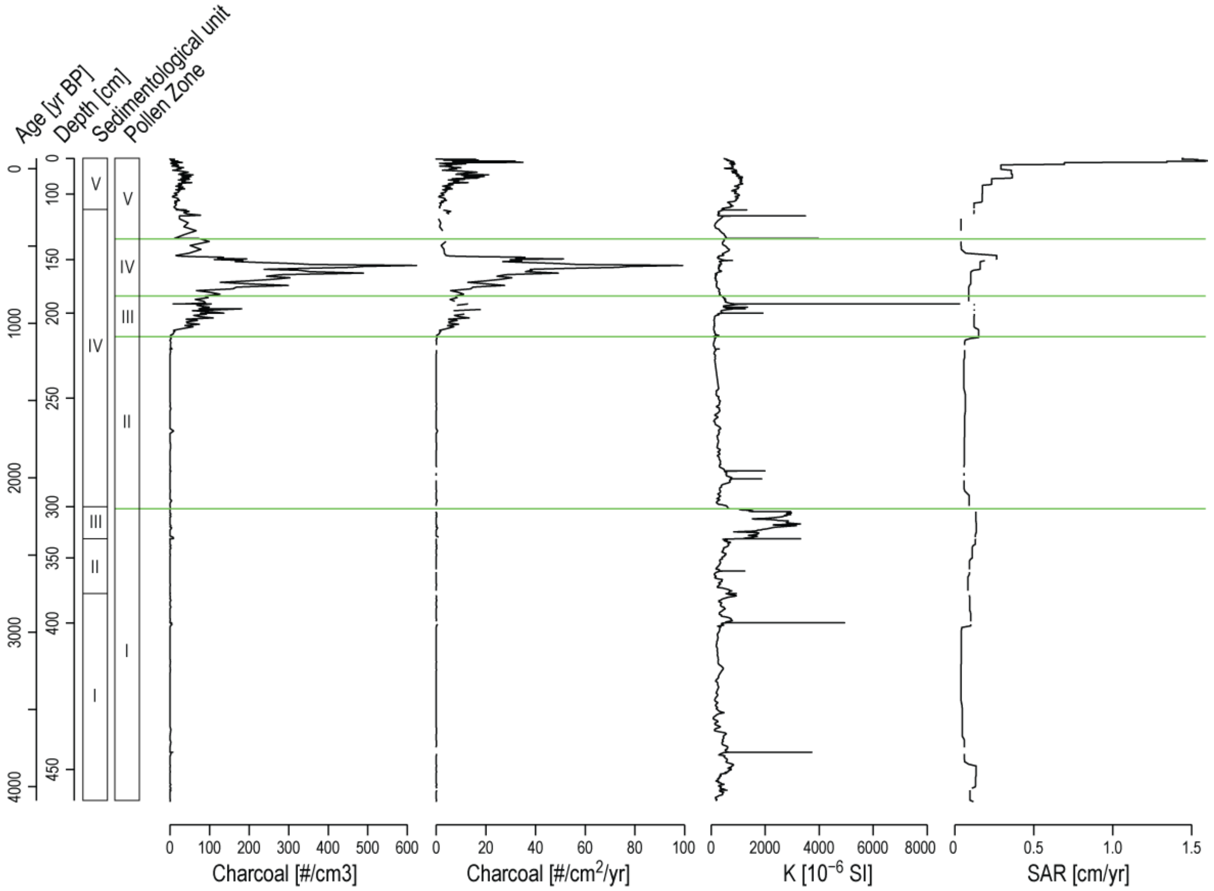


**Figure S4** Sedimentological units, pollen zones, concentration and influx of charcoal particles, magnetic susceptibility (K) and sediment accumulation rate (SAR) of the composite sediment core LMAHA18 plotted on age scale (indicated in calendar years before present) and on depth.

**Table S3**. Bibliographic review of the demographic studies currently available for Malagasy wildlife distributed across north of Madagascar using molecular datasets (n = 19 study species). WGS = Whole Genome Sequences; RADseq = Restriction-site Associated DNA sequencing; ddRADseq = double digestion Restriction-site Associated DNA sequencing; mtDNA = mitochondrial DNA; PSMC = Pairwise Sequentially Markovian Coalescent; BSP = Bayesian Skyline Plot; EBSP = Extended Bayesian Skyline Plot; ABC = Approximate Bayesian Computation framework; bot = bottleneck; kyr = thousand years.

| Species | Vernacular name | Taxa | Habitat type | Study area | Dataset | Method | Demographic event | Time of event (kyr) | Reference |
|---|---|---|---|---|---|---|---|---|---|
| *Microcebus arnholdi* | Arnhold's Mouse Lemur | mammal | humid | Montagne d'Ambre | WGS; RADseq | Stairway Plot | bottleneck, recover, bottleneck | > 25 (1° bot); < 5.5 kyr (2° bot) | [5] |
| | | | | | | PSMC | bottleneck, recover, bottleneck | 25 (1° bot); < 5.5 kyr (2° bot) | |
| | | | | | | fastsimcoal2 | Two bottlenecks | 1 and 5 | |
| *Microcebus tavaratra* | Northern Rufous Mouse Lemur | mammal | both | Loky – Manambato region | mtDNA (Dloop, cytb, cox2) | mismatch distribution, neutrality tests | expansion | – | [6] |
| | | | | | | EBSP | constant pop size | – | |
| *Propithecus perrieri* | Perrier's Sifaka | mammal | dry | Analamerana – Andrafiamena region | microsatellites | BOTTLENECK | bottleneck | – | [7] |
| *Propithecus perrieri* | Perrier's Sifaka | mammal | dry | Analamerana – Andrafiamena region | microsatellites | MSVAR | bottleneck | 3.3 – 10 | [8] |
| | | | | | | VAREFF | bottleneck | 6 – 45 | |
| | | | | | | ABC | bottleneck | 1 – 3.4 | |

| | | | | | | | | | |
|---|---|---|---|---|---|---|---|---|---|
| *Propithecus perrieri* | Perrier's Sifaka | mammal | dry | Analamerana – Andrafiamena region | microsatellites | BOTTLENECK | bottleneck | – | [9] |
| *Propithecus tattersalli* | golden-crowned sifaka<br>sifaka | mammal | both | Loky – Manambato region | microsatellites | BOTTLENECK | bottleneck | – | [9] |
| *Propithecus tattersalli* | golden-crowned sifaka<br>sifaka | mammal | both | Loky – Manambato region | microsatellites | VAREFF | bottleneck | 0.6 – 5.4 | [8] |
| | | | | | | ABC | bottleneck | 0.3 – 1.5 | |
| *Propithecus tattersalli* | golden-crowned sifaka<br>sifaka | mammal | both | Loky – Manambato region | microsatellites | MSVAR | bottleneck | 5.5 – 10 | [10] |
| *Eliurus tanala* | Tanala tufted-tailed rat | mammal | humid | Loky – Manambato region | mtDNA (cytB, D-loop) | mismatch distribution, neutrality tests | expansion | – | [11] |
| | | | | | | BSP | constant pop size | – | |
| *Eliurus carletoni* | Ankarana Special Reserve Tufted-tailed Rat | mammal | dry | Loky – Manambato region | mtDNA (cytB, D-loop) | mismatch distribution, neutrality tests | bottleneck | – | [11] |
| | | | | | | BSP | bottleneck | 15 | |
| *Eliurus carletoni* | Ankarana Special Reserve Tufted-tailed Rat | mammal | dry | Loky – Manambato region, Ankarana, Analamerana | mtDNA (cytB, D-loop) | neutrality tests Bayesian skyline plot | inconclusive | – | [12] |
| | | | | | | BSP | bottleneck | 7.5 – 18.75 | |
| *Microgale brevicaudata* | – | mammal | both | Montagne d'Ambre; Montagne des | mtDNA (ND2) | neutrality tests | expansion | – | [13] |

| | | | | | | | | | |
|---|---|---|---|---|---|---|---|---|---|
| | | | | Français; Loky – Manambato region | | BSP | expansion | – | |
| *Chaerephon leucogaster* | Grandidier's free-tailed bat | mammal | dry | Nosy Be, Nosy Komba, Ambilobe | mtDNA (cytB, D-loop) | mismatch distribution, neutrality tests | expansion | – | [14] |
| | | | | | | BSP | expansion; bottleneck | 29; < 4 | |
| *Chaerephon leucogaster* | Grandidier's free-tailed bat | mammal | dry | Nosy Be, Nosy Komba, Ambilobe | mtDNA (cytB, D-loop) | neutrality tests | expansion | 5.8 – 11.1 | [15] |
| *Chaerephon leucogaster* | Grandidier's free-tailed bat | mammal | dry | Antsiranana | mtDNA (cytB) | BSP | constant pop size | – | [16] |
| *Myotis goudoti* | Malagasy Mouse-eared Bat | mammal | both | North Madagascar | mtDNA (cytB, D-loop) | mismatch distribution, neutrality tests | expansion | 110.8 – 128.6 | [17] |
| *Dicrurus forficatus* | Crested Drongo | bird | both | Manongarivo Special Reserve | mtDNA (ND2, ATP6), autosomal loci | neutrality tests | expansion | LGM | [18] |
| *Ravenea robustior* | – | Palm tree | dry | Sava region | ddRADseq | EBSP | expansion | – | [19] |
| *Laliostoma labrosum* | Madagascar Bullfrog | amphibian | both | Ankarana Special Reserve | mtDNA (16S) | ABC | expansion | late Quaternary | [20] |
| *Calumma boettgeri* | – | non-avian reptile | both | Montagne d'Ambre | mtDNA (ND2) | ABC | expansion | late Quaternary | [20] |

| | | | | | | | | | |
|---|---|---|---|---|---|---|---|---|---|
| *Furcifer pardalis 1* | Panther Chameleon | non-avian reptile | humid | NW coast, Nosy Komba | mtDNA (cytB) | ABC | expansion | late Quaternary | [20] |
| *Furcifer pardalis 7* | Panther Chameleon | non-avian reptile | dry | Montagne d'Ambre | mtDNA (cytB) | ABC | expansion | late Quaternary | [20] |
| *Furcifer pardalis 3* | Panther Chameleon | non-avian reptile | humid | Nosy Be | mtDNA (cytB) | ABC | not expanding | late Quaternary | [20] |
| *Furcifer pardalis 8* | Panther Chameleon | non-avian reptile | dry | Antsiranana | mtDNA (cytB) | ABC | not expanding | late Quaternary | [20] |
| *Geckolepis maculata* | Fish-scale Gecko | non-avian reptile | both | Montagne d'Ambre, Montagne des Français | mtDNA (ND4) | ABC | not expanding | late Quaternary | [20] |
| *Madagascarop his colubrinus* | – | non-avian reptile | dry | Antsiranana | mtDNA (COI) | ABC | expansion | late Quaternary | [20] |
| *Mimophis occultus* | Northern Big-eyed Snake | non-avian reptile | both | Ankarana Special Reserve, Ambanja, Antsiranana | mtDNA (COI) | ABC | expansion | late Quaternary | [20] |
| *Zonosaurus madagascarien sis* | – | non-avian reptile | both | Loky – Manambato region | mtDNA (cytB) | ABC | not expanding | late Quaternary | [20] |
| *Phelsuma dorsivittata* | – | non-avian reptile | humid | Montagne d'Ambre | mtDNA (16S) | mismatch distribution, neutrality tests | bottleneck | – | [21] |
| | | | | | | BSP | bottleneck | – | |
